# SifA SUMOylation governs *Salmonella* Typhimurium intracellular survival via modulation of lysosomal function

**DOI:** 10.1101/2023.03.02.530793

**Authors:** Hridya Chandrasekhar, Gayatree Mohapatra, Mukesh Singh, Sarika Rana, Navneet Kaur, Sheetal Sharma, Amit Tuli, Prasenjit Das, C. V. Srikanth

## Abstract

Gastroenteritis causing pathogen *Salmonella* Typhimurium (*S.* Tm) during its infection in host cells thrives in a vacuolated compartment, *Salmonella* Containing Vacuole (SCV), which sequentially acquires host endosomal and lysosomal markers. Long tubular structures, called as *Salmonella* induced filaments (SIFs), are known to be required for SCV’s nutrient acquisition, membrane maintenance and stability. A tightly coordinated interactions involving prominent effector SifA and various host adapters PLEKHM1, PLEKHM2 and Rab GTPases govern SCV integrity and SIF formation. Here, we report for the first time, the functional regulation of SifA is modulated by its SUMOylation at lysine 11. *S.* Tm expressing lysine 11 mutant SifA (SifA^K11R^) is defective in intracellular proliferation due to compromised SIF formation and enhanced lysosomal acidification. Furthermore, murine competitive index experiments reveal defective in vivo proliferation and weakened virulence of SifA^K11R^ mutant. Concisely, our results demonstrate that SUMO deficient SifA mutant nearly behaves like a SifA knockout strain which impacts PLEKHM2-M6PR mediated lysosomal acidification pathway. Thus, our results bring forth a novel *S.* Tm-host crosstalk mechanism involving host mediated effector SUMOylation critical for pathogenicity.

## Introduction

The Gram-negative bacterium *Salmonella* Typhimurium (hereafter to be referred as *S*. Typhimurium or *S*. Tm) causes self-resolving gastroenteritis in humans. Although, the infection clears out in a week’s span in healthy individuals, there is impending danger of bacteremia in immunocompromised individuals and infants. Even in the current scenario, infections caused by *S*. Tm are frequent, leading to a significant health burden in both developing and developed world Zhang *et al*., 2003).

*S.* Tm pathogenicity is attributed to the presence of unique cluster of genes in its genomes making up the *Salmonella* pathogenicity islands. Among many SPIs, *Salmonella* pathogenicity island 1 (SPI1) and *Salmonella* pathogenicity island 2 (SPI2) are well explored (Marcus *et al*., 2000). Encoded within these SPI islands are a bunch of effectors with diverse functions, their associated chaperones, and proteins structuring Type III Secretion System (Deng *et al*., 2017) assisting the delivery of bacterial effectors in the host. A coordinated action of SPI-1 effector proteins aid in *S.* Tm entry into host along with induction of proinflammatory signaling mechanisms (Marcus *et al*., 2000; McGhie, Hayward and Koronakis, 2004; Lou *et al*., 2019).

Post uptake *S.* Tm reside in a secluded membranous compartment called *Salmonella* Containing Vacuole (SCV) which sequentially acquires markers of endocytic pathway. Early Endosome Antigen 1 (EEA1), GTPases Rab5 and Rab11 are initially recruited to SCV which are later interchanged with late endocytic markers Rab7, Lysosomal Associated Membrane Proteins 1,2.3 (LAMPs) and vacuolar ATPases (Steele-Mortimer, 2008). Sequestration of *S.* Tm within SCV is important since it enables intracellular replication and evasion of host defense mechanisms (Birmingham and Brumell, 2006; Birmingham *et al*., 2006). Members of the Rab family are crucial regulators of the intracellular membrane trafficking, thereby, they also modulate the fate of SCV (Smith *et al*., 2007). Aided by effector functions, *S.* Tm are known to target Rab7 to modify SCV maturation (Méresse *et al*., 1999). Recently our group also showed that *S.* Tm manipulates Rab7 function by interfering with its SUMOylation, a post-translational modification mechanism (Mohapatra et al., 2019). Several crucial aspects of SCV biogenesis and its stability are majorly fine-tuned by an SPI-2 effector *Salmonella* induced filament A (SifA) (Beuzón *et al*., 2000). SifA directs PLEKHM1 (Pleckstrin homology and RUN domain containing M1), Rab7 and HOPS (Homotypic fusion and protein sorting) complex to mobilize late endosomal membrane compartments for SCV generation (McEwan *et al*., 2015). PLEKHM1 directly interacts with both Rab7 and SifA, through its Pleckstrin homology domain (PH domain) (Rawet-Slobodkin and Elazar, 2015). A ΔSifA mutant of *Salmonella* escapes SCV and fails to replicate in macrophages and thereby ends in a compromised pathogenicity *in vivo* (Brumell, Rosenberger, *et al*., 2001).

Tube like membranous extensions generated from SCV membrane, collectively called as *Salmonella* induced filaments (SIFs), are generated during later stages of infection (Drecktrah *et al*., 2008; Rajashekar *et al*., 2008). SIFs are required for SCV nutrition and stability (Kuhle, Abrahams and Hensel, 2006; Zhang and Hensel, 2013; Liss *et al*., 2017). SifA function engages microtubule dynamics which is required for SIF formation (Brumell, Goosney and Finlay, 2002) as well as recruitment of lysosome associated membrane protein 1 (LAMP1) on to the tubules (Garcia-del Portillo *et al*., 1993; Stein *et al*., 1996). Furthermore, through SifA dependent mechanisms, *S.* Tm targets PLEKHM1 and SKIP (PLEKHM2) (Dumont *et al*., 2010). SifA forms a complex with SKIP which enable regulation of plus-end-end directed motor kinesin-1 on SIFs (Diacovich *et al*., 2009a; Fang *et al*., 2022) (Boucrot *et al*., 2005). Though, SCV later fuses with lysosomes, they are devoid of recycling mannose-6-phosphate receptors (MPRs) responsible for trafficking lysosomal hydrolases into lysosomes (Garcia-del Portillo and Finlay, 1995; Rathman, Barker and Falkow, 1997). The GTPase Rab9 through its interaction with SKIP associates with MPRs for routing these hydrolases into lysosomes (Barbero, Bittova and Pfeffer, 2002; McGourty *et al*., 2012). SifA and Kinesin interacting protein (SKIP) antagonistically prevents Rab9-SKIP association (Jackson *et al*., 2008). In this way, Rab9 dependent MPR pathway is inhibited, which in turn leads to blockade of lysosomal acidification (McGourty *et al*., 2012). Furthermore, impairment of RILP recruitment to Rab7, required for *S*. Tm survival, occurs through SifA function (Harrison *et al*., 2004). Thus, several mechanisms and functions crucial to Salmonellosis are attributed to SifA via its interaction with numerous host adapters. Intriguingly, SifA does not physically interact with Rab7, a host protein involved in the complex. Being able to carry out several regulatory roles by SifA is a paradigm, the mechanistic details of which is still not fully understood. To accomplish multiple functions, effectors often manipulate host PTMs.

Our group demonstrated that *S.* Tm control several aspects of host cell by altering its SUMOylation machinery (Verma *et al*., 2015). SUMOylation, a small ubiquitin like modifier systems which is known to post-translationally modify proteins and thereby affecting their function. SUMOylation being a powerful machinery can impact stability, folding, interaction and localization of their substrates (Wilkinson and Henley, 2010). The global SUMOylome in hosts undergoes a major alteration during *S*. Tm infection, several of which belong to vesicular transport pathway (Mohapatra et al., 2019). Also, bacterial effectors from certain pathogens *Anaplasma* (Beyer *et al*., 2015) and *E. chaffenis* (Dunphy, Luo and McBride, 2014) undergoing SUMOylation has been shown to be beneficial for pathogenesis. It is unclear if SifA or any of *S.* Tm effectors undergo PTM modification by SUMO. This study aims to identify whether SifA undergoes such processes and if so, how this modification could add complexity to *S.* Tm pathogenicity.

## Results

### Lysine 11 of SifA gets SUMOylated during infection

Since, SifA and Rab7 reside in proximity in the SCV and Rab7 SUMOylation is prevented by *S.* Tm during infection (Mohapatra *et al*., 2019), we investigated possible interaction of SifA with components of SUMOylation machinery. In our preliminary analysis, a yeast two hybrid assay was conducted to probe the interaction between SifA with Smt3, a yeast SUMO1 ortholog or Ubc9, the human E2 enzyme of SUMOylation pathway. The genes encoding SifA, Smt3, Smt3ΔGG and human Ubc9 were cloned in both yeast bait expression vector pGBKT7 expressing GAL4 DNA binding domain (BD) and pGADT7 vector designed to express GAL4 activation domain (AD) separately. Yeast cells (Y187, Clonetech’s matchmaker Y2H system) expressing both bait (pGBKT7) and prey vector (pGADT7) were confirmed by their growth on selection plates (Fig 1a). We observed that both Smt3 and Ubc9 showed interaction with SifA as seen by the growth of yeast on plates devoid of Histidine and Adenine. Notably, the mutant protein Smt3ΔGG, where C-terminal di glycine motif was deleted showed no growth. Screening was further carried on selection plates containing 2-Amino-1H-1,2,4-triazole (3-AT) at a concentration of 2.5mM adding stringency to the histidine-based growth selection. Similarly, selection comprising either one empty parent vector or between parent vectors led to complete inhibition of growth (fig 1a). These data led us to conclude that SifA interacts with Smt3 and Ubc9. It was therefore important to examine if this interaction was a result of SifA undergoing SUMOylation.

**Fig. 1:**
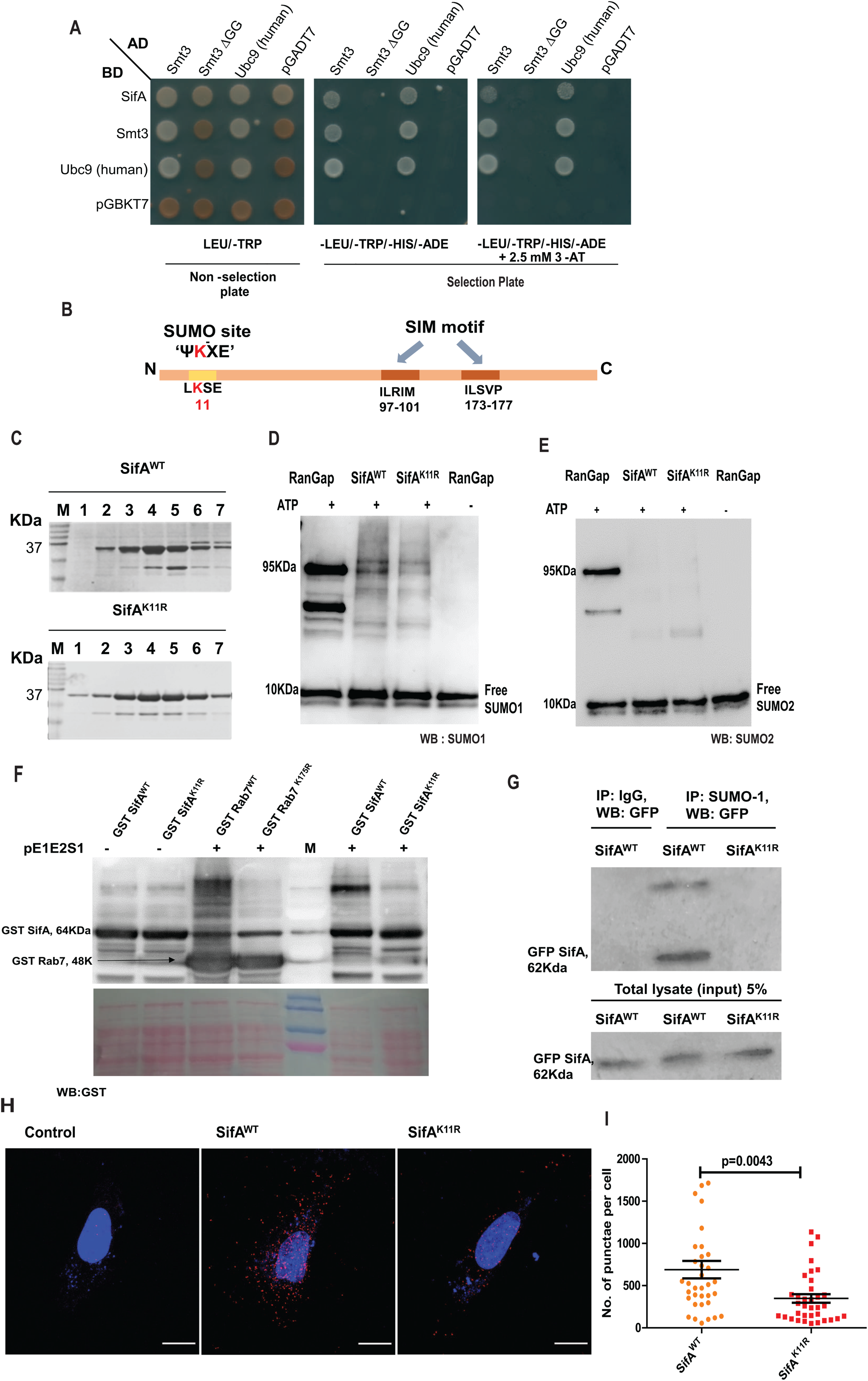
Lysine 11 of SifA gets SUMOylated during infection. **(a)** Y2H assay indicating interaction between SifA and SUMO1, Smt3 (yeast SUMO1 ortholog), Smt3ΔGG (Smt3 with C-terminal di glycine deleted) and human Ubc9 (E2 enzyme of SUMO pathway). **(b)** Schematic representation of SifA primary sequence highlighting SUMO consensus motif and SUMO interacting motifs (SIM). The lysine at 11^th^ position within a SUMO consensus site was predicted as a potential SUMOylation site with high probability by multiple in silico tools. **(c)** Fractions of purified GST-SifA^WT^ and GST-SifA^K11R^ obtained from superdox 200 column run on an SDS PAGE (12%) and Coomassie stained for expression analysis. **(d, e)** Resultant immunoblot of isolated proteins subjected to *in vitro* SUMOylation assay separately probed for both SUMO1 and SUMO2/3 isoforms. The results indicated the presence of higher molecular bands in wild type SifA incubated with SUMO1 isoform, whereas the intensity of bands was remarkably less in K11R mutant. **(f)** In-bacto SUMO conjugation assay for GST tagged SifA expressed in *E. coli* BL21 along with pE1E2S1 plasmid encoding SUMO enzyme machinery. The blots were probed by anti-GST antibodies. Rab7 was used as a positive control (*Mohapatra et al*). **(g)** Co-immunoprecipitation of eGFP SifA expressed in HCT-8 cells pulled down by SUMO-1 antibody, succeeding immunoblot probed by anti-GFP antibody showed interaction only in wild-type SifA expressing lysates. **(h)** Images displaying discreet PLA punctae representing colocalization of SifA and SUMO-1 in indicated samples (scale 6 micron). **(i)** Quantification of PLA punctae carried out using particle analysis tool from Image J software.

The primary sequence of SifA was analyzed through various online ‘SUMO’ prediction software, including GPS-SUMO, JASSA and SUMO plot analysis program were employed for this purpose (Zhao *et al*., 2014; Beauclair *et al*., 2015), for identifying the presence of probable SUMO consensus sites. The search from all these above-mentioned programs collectively pointed at lysine11 (K11) lying within a SUMO consensus motif ΨKXE/D (Sampson, Wang and Matunis, 2001) displaying high score in comparison to the cut-off score (Table 1). Apart from the indicated SUMO consensus sites, primary sequence of SifA accommodate two probable SIM sites spanning from amino acids 97-101 and 173-177 as depicted in (fig1b).

**Table 1.** Table displaying potential SUMOylatable lysines within SUMO consensus sites and SIM sites from primary sequence of SifA as predicted by indicated software.

| Software | SUMO site identified (Lysine) within SUMO consensus (Ψ-K-X-E) | Score | SIM site (V/I-X-V/I-V/I) | Score |
| --- | --- | --- | --- | --- |
| GPS-SUMO | 11 | 14.818 | 97-101,<br>173-177 | 29.92 |
| Agilent SUMOplot | 11, 35 | 0.91 (high),<br>0.59 | NIL |  |

To assess the possibility of SUMOylation at lysine 11, a mutant was generated with an arginine substitution (K11R) via site directed mutagenesis. The recombinant proteins wildtype SifA GST-SifA^WT^ and lysine mutant (GST-SifA^K11R^) were expressed in *E. coli* Rosetta strain and purified (fig 1c). *In vitro* SUMOylation assay was conducted on these purified fractions of SifA, using components from ENZO SUMOylation kit. Post reaction the mixture was run on SDS PAGE and probed with anti-SUMO-1 and anti-SUMO2/3 antibody separately. As indicated in the immunoblot in fig 1d, higher molecular weight bands were observed in SifA^WT^ when probed with SUMO1 antibody, whereas the intensity of bands were significantly less in SifA^K11R^. Moreover, lack of any detectable band from the immunoblot probed by anti-SUMO2 suggests possible SUMO1-modification of SifA at lysine 11 (fig 1e). Appearance of higher molecular weight bands in RanGAP1 (positive control) and clear lanes in reactions without ATP evinced the quality of SUMOylation reactions (1d, e).

The above observations were further reinforced by another assay referred to as ‘in bacto-SUMOylation assay.’ This assay majorly relies on the expression of plasmid pE1E2S1 encoding SUMO machinery enzyme components (E1, E2) and SUMO1 (S1) besides the expression of target protein, SifA in *E. coli* (Uchimura *et al*., 2004). The pE1E2S1 plasmid along with constructs encoding SifA^WT^ or SifA^K11R^ were co-transformed in *E. coli* BL21 cells. The lysates from these cells were run on SDS-PAGE and immunoblotted. As depicted in fig 1f, a prominent band around 120KDa was observed, indicating possible SUMO-modificartion of SifA, only in sample comprising co-transformed wild type SifA and pEIE2S1 construct, but not with those with pE1E2S1 and K11R mutant. The mammalian protein Rab7 and its corresponding SUMO mutant, Rab^K175R^ were also included here as a positive and negative control for the assay respectively (Mohapatra *et al*., 2019) (fig 1f). As anticipated, higher mobility bands in case of Rab7 but not for Rab^K175R^ was observed.

In order to validate these observations *in vivo* in mammalian cells, a co-immunoprecipitation (co-Ip) assay was conducted using human intestinal epithelial cell line, HCT-8. HCT-8 cells were transfected by plasmids bearing both SifA^WT^ and SifA ^K11R^ genes cloned in mammalian peGFP-C1 vector separately. After 24 hours of transfection, the transfected cells were subjected to infection by ΔsifA mutant for 7 hours. Immunoprecipitation was carried by anti-SUMO1 antibody followed by immunobloting with anti-GFP antibody. A band corresponding to eGFP SifA^WT^ around 62KDa and prominent upper band just in SifA wild type transfected samples suggesting interaction between SUMO1 and SifA was observed. Notably, in line with our above results, we did not observe any bands in SifA^K11R^ transfected samples making it indistinguishable from isotype control (fig 1g). The expression of SifA^K11R^ protein was observable in the input sample (fig 1g).

Finally, an in-situ detection of SUMO modified SifA in cells were accomplished through a proximity ligation assay (PLA). This assay was conducted in HeLa cells transfected by peGFPC1 plasmids containing SifA^WT^ or the non-SUMOylatable -K11R mutant SifA (hereafter to be referred as SUMO mutant SifA or SifA^SMUT^) for 24 hours. The transfected cells were probed by anti-GFP antibody (rabbit) and anti-SUMO-1 antibody (mouse). PLA mainly relies on the attachments of oligonucleotide labelled secondary antibodies against the primary that will only be joined and ligated if the interacting proteins are in proximity. The interaction is expected to result in the formation of a closed circular DNA template which is eventually amplified and detected by detection oligos which can be visualized as discreet puncta (fig 1g). In the control sample, one primary antibody (anti-GFP) was omitted. As shown in the fig 1h the puncta from individual cells quantified suggested interaction between SifA and SUMO-1. The number of punctae was observed highest in SifA^WT^ transfected cells in comparison to those with SifA^SMUT^ (fig 1i). While, there was no signal in the negative control (left panel). Based on the results from all these above experiments, we concluded that SifA undergoes SUMO-modification by SUMO-1 isoform at lysine 11.

### Ectopically expressed SifA ^SMUT^ displays a compromised stability

As it is well established that SUMOylation can potentially impact its substrate’s stability, folding, interaction or/and compartmentalization (Wilkinson and Henley, 2010), we therefore set out to investigate the biological consequences of SifA SUMOylation.

HCT-8 cells transfected by peGFPC1 plasmids containing SifA^WT^ or SifA^SMUT^ were monitored for indicated time periods as shown in fig 2a. There were no notable differences in expression at 24 and 36 hrs post transfection between SifA^WT^ and SifA^SMUT^. However, at 48 hrs post transfection, compared to SifA^WT^ a significant decrease in pools of SifA^SMUT^ was observed as shown in fig 2a, b. Next, cycloheximide chase assay was executed to determine the steady state stability and half-life of both proteins (Kao *et al*., 2015). To accomplish this, cycloheximide at a concentration of 100µg/ml was incorporated into transfected HCT-8 cells succeeding 24 hrs of transfection (fig 2c). The decay curve represents the mean of remaining percent protein acquired from three independent sets at each time point in comparison to start of the chase (time 0). As it is evident from the curve (fig 2d), the stability of SifA^K11R^ was considerably less compared to the wild type protein beginning at 2 hrs. At 4 hrs the pools of SifA^K11R^ were less than 25 % of the initial, whereas at this the SifA^WT^ around 50% of initial. Now to ascertain the pathway leading to the degradation of SifA^SMUT^ transfected cells were treated with MG132 (inhibitor of ubiquitin proteasomal pathway) or bafilomycin A1 (blocker of lysosomal degradation). It was evident that the levels of both the proteins did not alter significantly upon MG132 treatment (fig 2e, f), only a mild increase in SifA^SMUT^ seen. However, treatment of bafilomycinA1 restored the levels of SifA^SMUT^ (fig 2g, h). Thus, the relative pools of wild type and SifA^SMUT^ were identical in presence of this drug indicating a role of lysosomal activity in regulation SifA^K11R^ expression. Next, live cell imaging was carried out in SifA^WT^ or SifA^SMUT^ transfected HCT-8 cells, to assess the colocalization between SifA and lysosomes. Lysosomes were stained by the cell permeable fluorescent dye lysotracker red. Live cell snaps acquired via confocal imaging displayed diminished GFP fluorescence (green channel) in case of GFP SifA^SMUT^ affecting its colocalization with lysosomes in contrast to GFP SifA^WT^ which could exquisitely associate with lysosomes (fig 2i). Hence, the data from above experiments implied that ectopically expressed SifA^SMUT^ is more prone to lysosomal degradation compared to its wildtype counterpart.

**Fig. 2:**
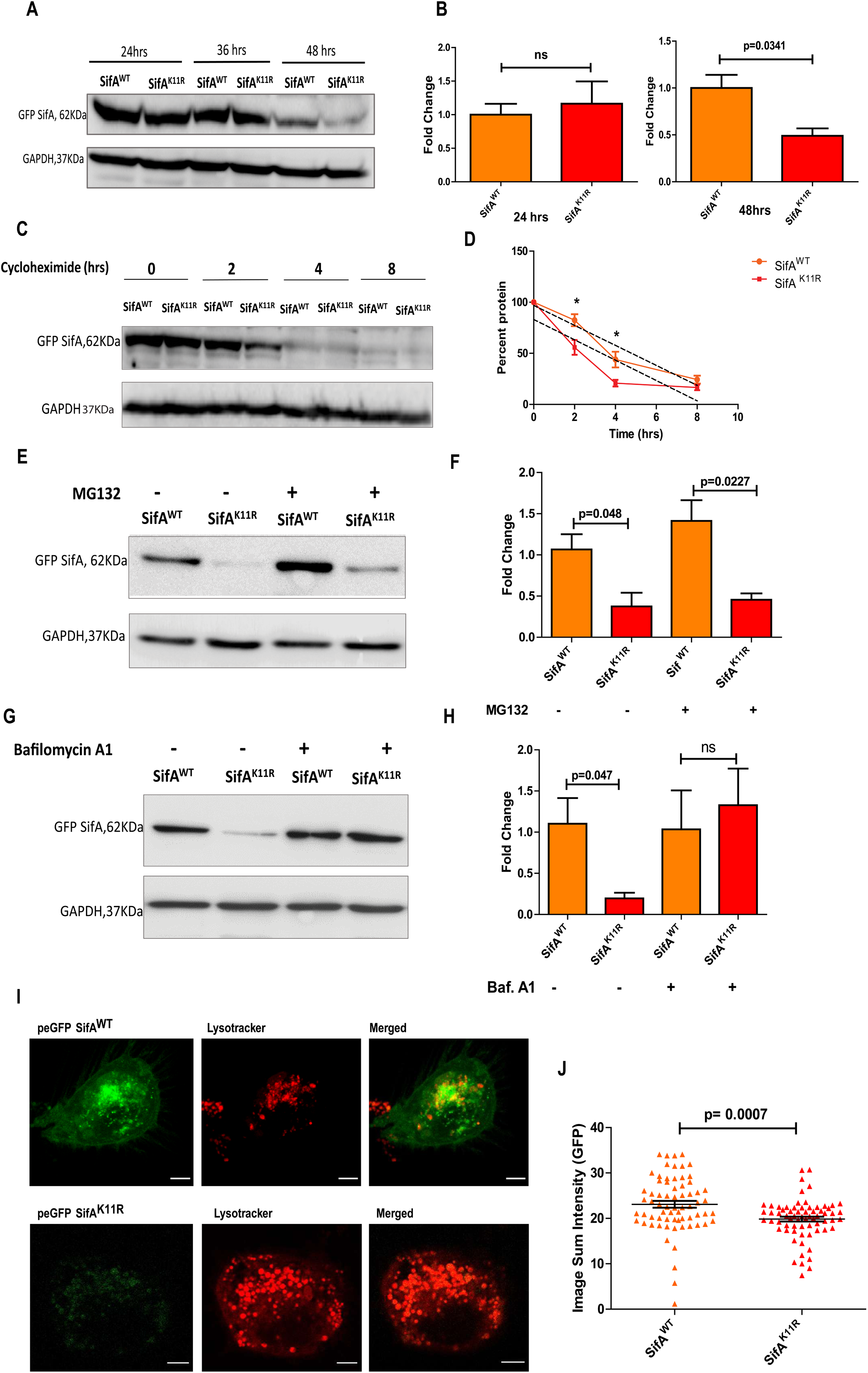
Ectopically expressed SifA ^K11R^ (SMUT) protein is unstable in comparison to wild-type. **(a)** Timeline of expression levels of ectopically expressed SifA^WT^ and SifA^SMUT^ proteins from HCT-8 cell lysates. A drastic difference in the levels were detected after 48 hrs of transfection. **(b)** The normalized levels of SifA^SMUT^ are represented in comparison to expression levels of SifA ^WT^ protein taken as the control for indicated time periods. **(c)** Cycloheximide (100µg/ml) was incorporated into SifA^WT^ and SifA^SMUT^ transfected HCT-8 cells after 24 hrs of transfection. Expression levels of both proteins are indicated till 8 hours of treatment duration (cyclo chase). **(d)** Decay curve indicating the stability of both proteins. The level of each protein at time 0 was set as 100% and the percentage of protein remaining at each time point from three independent sets was calculated and plotted (p value corresponds to 0.0457 at 2hrs and 0.0368 at 4hrs). **(e)** Blots representing the expression levels of wild-type and SUMO mutant SifA and its relative quantification **(f)** on proteasomal inhibitor MG132(20µM) treatment**. (g, h)** Impact of lysosomal acidification inhibitor bafilomycin A1 (100nM) on the expression alteration of SifA ^SMUT^ in comparison to the wildtype protein. **(i)** Live cell snapshots indicating localisation of SifA wild type and SMUT with lysosomes labelled by lysotracker red (scale 5 microns).

### SUMOylation of SifA is required for preventing lysosomal acidification and eventual lysosomal clearance of *Salmonella*

Taking into consideration, the role of lysosomes in regulating SifA’s stability from our previous findings and the fact that one of major functions of this effector is protection of the bacterium from lysosomal degradation, we decided to decode how each downstream adapters involved in this process respond to SifA SUMOylation. As mentioned in the introduction, SifA is involved in subverting retrograde trafficking of mannose-6-phosphate receptors (MPRs) hence inhibiting lysosomal acidification. It is known that SifA-SKIP complex is formed during *Salmonella* infection enables sequestration of Rab9 protein leading to misrouting of MPRs (McGourty *et al*., 2012). Also, the host interactor of SifA, PLEKHM2 (SKIP) is regarded as a negative regulator of lysosomal activity (McGourty *et al*., 2012) and recently its expression was known to be modulated by certain strain of *Mycobacteria* (Laopanupong *et al*., 2021). We began by examining whether SKIP expression gets modulated during *Salmonella* infection. Accordingly, HCT-8 cells were infected by the following strains of *Salmonella*; SL1344 (wild type), ΔSifA (SifA gene knock-out in SL1344 background), ΔSifA complemented with SifA^WT^ (pSifA^WT^) or SifA^SMUT^ (pSifA^SMUT^) (cloned in pBH-HA vector) individually at a MOI of 1:40 for 4 hrs. Samples infected by SL1344 showed a slight increase in SKIP levels compared to lysates generated from uninfected control, but the levels did not rise in ΔSifA infected samples (fig 3a). Amidst pSifA^WT^ and pSifA^K11R^ infected samples, collected at 4hpi, there was a considerable difference in SKIP expression (p value=0.056) (fig 3a) with a sizeable reduction in SKIP levels in pSifA^SMUT^ infected samples. Lysates obtained from ectopically expressed SifA^WT^ and SifA^SMUT^ in HCT-8 cells followed by ΔSifA infection (4hrs, MOI 1:40) after 24 hrs of transfection exhibited a drastic variation in the expression of protein Cation Independent mannose 6 phosphate receptor (CI-M6PR) which was unforeseen (fig 3 b). The levels of CI-M6PR protein were enhanced in case of SifA^SMUT^ expressing lysates. We surmised that SifA^SMUT^ is deficient in the known function of preventing lysosomal acidification like its wild type counterpart. To pursue this, the ability of Rab9 to colocalize with CI-M6PRs was compared during an infection with complemented strains pSifA^WT^ and pSifA^SMUT^. Colocalization assays were conducted in HeLa cells, as this cell line is widely employed for imaging studies related to *Salmonella* infection (Giannella *et al*., 1973; Niesel, Chambers and Stockman, 1985; Misselwitz *et al*., 2010). HeLa cells were infected by pSifA^WT^ or pSifA^SMUT^ separately at a MOI of 1:40 for a duration of 7 hours, after which these infected cells were fixed, stained for CI-M6PR (green channel), Rab9 (red channel) cell nuclei and indicated strains were highlighted by DAPI (blue channel) (fig 3c). The analysis from more than ninety cells represented by individual dots portrayed a statistically significant increase in colocalization between CI-M6PRs and Rab9 in case of cells infected by pSifA^SMUT^ compared to wild-type (fig 3d).

**Fig. 3:**
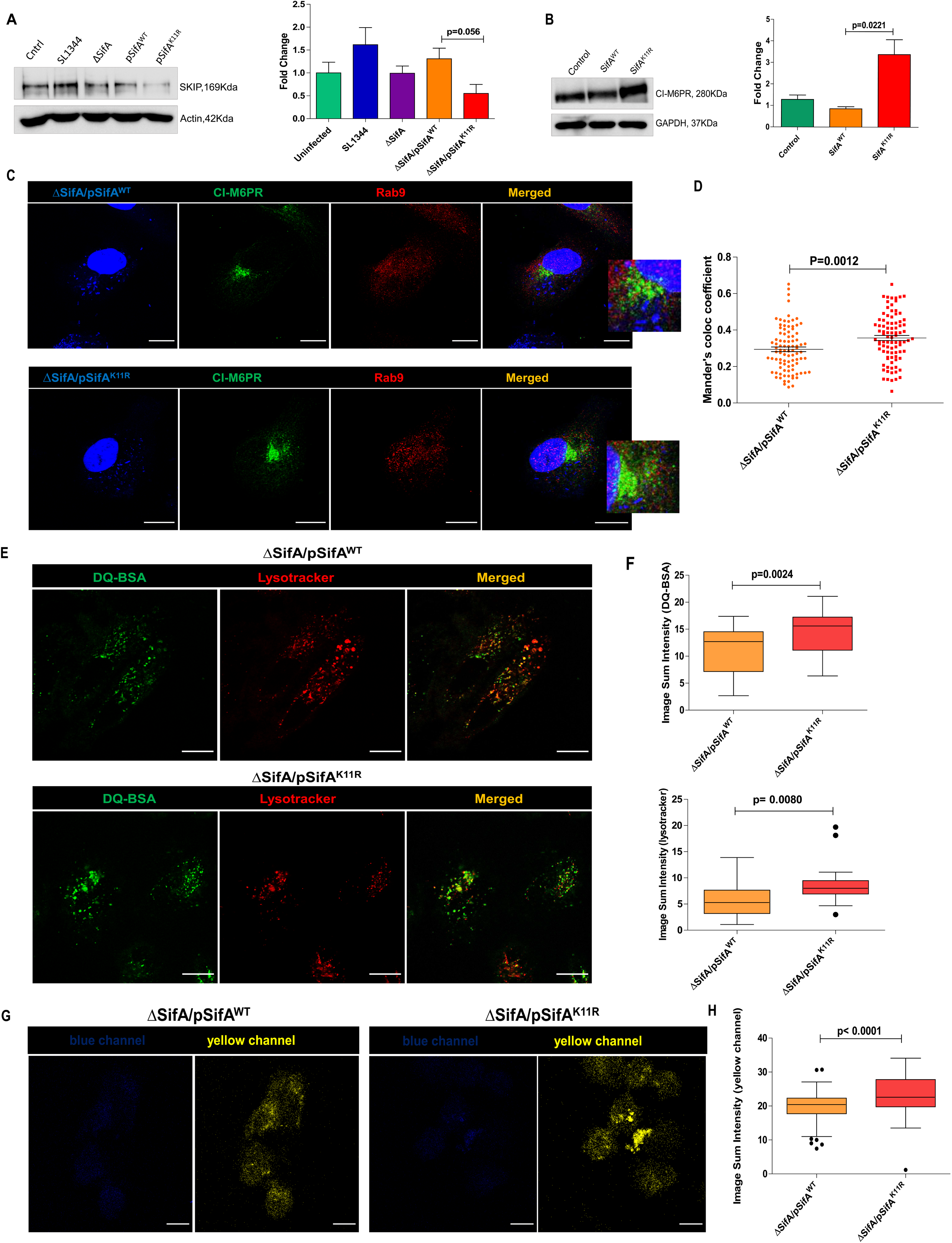
SUMOylation of SifA is required for preventing lysosomal acidification and eventual lysosomal clearance of *Salmonella*. **(a)** PLEKHM2 (SKIP) expression levels and their quantification in response to infection with depicted strains of *Salmonella*. **(b)** Immunoblot and its densitometric analysis representing CI-M6PR expression in transfected samples followed by ΔSifA infection. Control refers to lysates prepared from non-transfected samples. **(c)** Images stained for Rab9 and CI-M6PR for co-localization analysis in HeLa cells under pSifA^WT^ (scale 9micron) and pSifA^SMUT^ (scale 11 microns) infection conditions. **(d)** Analysis of Rab9 and CI-M6PR co-localisation comparison between pSifA^WT^ and pSifA^SMUT^ infected cells performed using Image J software. **(e)** Images of HeLa cell fields infected by shown strains acquired for lysosomal activity measurement using the DQ-BSA green probe, which emits bright green fluorescence in response to acidified environments (scale 14 micron). **(f)** Mean intensity of DQ-BSA green probe from images quantified using image-J software. **(g)** Lysosomal acidification was further monitored in infected cells stained by lysosensor yellow/blue DND. The yellow fluorescence (acidic environment) emitted by this ratiometric probe indicates the lysosomal acidity (scale 25micron). **(h)** Quantification and comparison of yellow fluorescence (acidic environment) emitted from lysosensor stained cells across indicated samples.

The inference from above observations concur SUMOylation of SifA at 11^th^ lysine is indispensable for overturning Rab9 MPRs trafficking. Thus, it is evident that SifA^SMUT^ is incapable of providing protection to SCV from digestion by lysosomal hydrolases. The predicted outcome in the current scenario stipulates escalated lysosomal activity in pSifA^SMUT^ infected cells compared to wild type SifA (pSifA^WT^).

Next, HeLa cells were incorporated with DQ-BSA green probe alongside indicated strains for a total duration of 7 hours. DQ-BSA is fundamentally made of BSA conjugated to a fluorogenic substrate, that are readily taken up by endocytosis and upon hydrolysis by lysosomal enzymes yield a bright green fluorescence (Frost *et al*., 2017; Marwaha and Sharma, 2017). To validate the green fluorescence, the dye lysotracker red was also added to the cells for the last 1 hour of infection, after which they were imaged without fixing. As displayed in fig 3e, the upper panel represents images obtained from pSifA^WT^ infected cells whereas the lower part shows images of cells bearing pSifA^SMUT^ infection. Colocalization between green and red channel indicated the green fluorescence is indeed emitted from lysosomal compartments. The intensity of both green and red channels emitted from pSifA^SMUT^ infected samples was found to be significantly high in collation to that of pSifA^WT^ infected cells (fig 3f). In addition to DQ-BSA, a similar ratiometric probe lysosensor yellow/blue DND was used to monitor the difference in lysosomal acidity across the samples. This dye shifts from blue to bright yellow as the pH shifts from neutral to acidic environment (DePedro and Urayama, 2009; Albrecht, Tejeda-Muñoz and De Robertis, 2020). As shown, from the fields corresponding to infection with pSifA^WT^ and pSifA^SMUT^ the intensity of yellow panel was several folds higher in latter compared to the former, thus implying that SifA^SMUT^ is incapable of lowering the lysosomal hydrolase activity (fig 3g, h). From here on, we concluded that SUMOylation of SifA is necessary for *Salmonella* to overcome degradation by lysosomal hydrolases.

### Cells infected by SMUT SifA (**SifA^SMUT^**) displays fewer SIFs

As mentioned earlier, *S*. Tm pathogenicity relies greatly on the its ability to generate SCV and SIFs in the endocytosed host cells, both of which are majorly attributed to effector SifA (Beuzón *et al*., 2000; Brumell, Goosney and Finlay, 2002). These phenotypes were surveyed in cells under SifA^SMUT^ infection to understand how SUMOylation regulate these functions of SifA. The protein galectin-8 is regarded as an indicator of SCV integrity. Galectin-8 binds to exposed host glycans on damaged SCVs activating xenophagy in host (Thurston *et al*., 2012). HeLa cells infected by complemented strains possessing SifA^WT^ and SifA^SMUT^ at an MOI of 1:40 for 4hrs were processed for immunofluorescence, to determine the acquisition of galectin-8 protein on SCV. From fig 4a it was evident that gal-8 (green channel) localized on SCV (red channel) in similar fashion during infection by both pSifA^WT^ and pSifA^SMUT^ strains which is also represented graphically in fig 4b. Proceeding further, infected HeLa cells were enumerated for visible SIFs to understand the effect of SifA SUMOylation on SIF generation. In general SIFs are formed at later hours of infection by fusion of SCV with late endocytic compartments carrying membranes enriched in glycoprotein LAMP1 (Brumell, Tang, *et al*., 2001). Thus, to enable visualization of SIFs, HeLa cells after 10hpi, were fixed and stained for the glycoprotein marker LAMP1. Image fields displaying LAMP1 (green channel) captured from cells infected by indicated strains are shown in fig 4c. The white arrows specify SIFs which appear like thin strands of thread. A zoomed image of single cell, each from infection by labelled pSifA^WT^ and pSifA^SMUT^ exhibiting SIFs were also shown in fig 4d. The number of SIFs counted from acquired frames corresponding to each strain were calculated in percentage and presented graphically in fig 4e. Overall, from the data it was clear that presence of SIFs was maximally seen in cells infected by bacteria expressing SifA^WT^. Besides no SIFs were produced in cells in response to ΔSifA infection. Intriguingly, no. of SIFs acquired from cells infected by bacteria expressing SifA^SMUT^ were substantially fewer compared to the wildtype strains, making it equivalent to ΔSifA. In view of the above data, we concluded that SUMOylation of SifA functionally regulates critical aspects related to *S.* Tm pathogenesis. We therefore set forth to investigate the survival and virulence potential of strain pSifA^SMUT^ in various infection models.

**Fig. 4:**
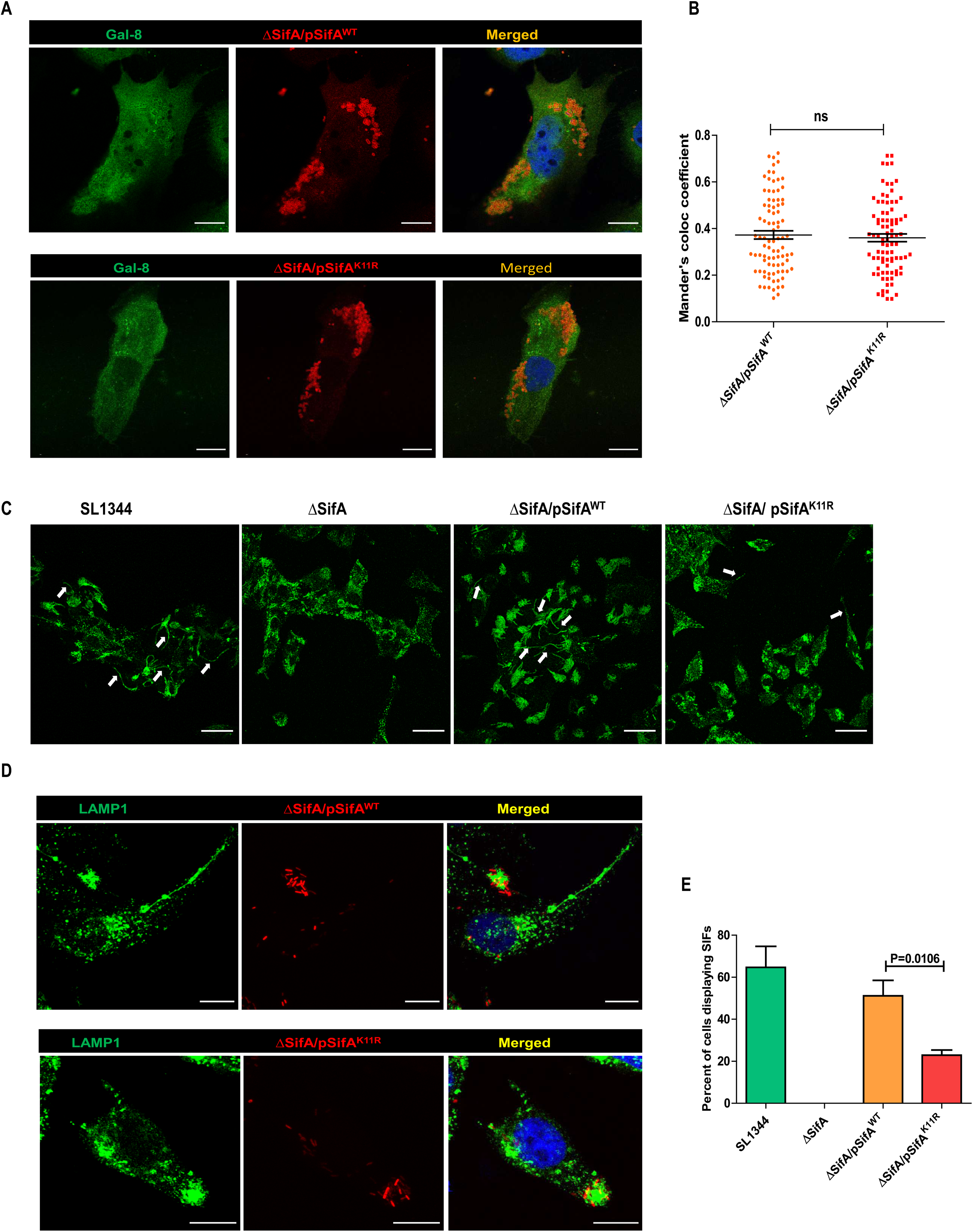
Cells infected by SMUT SifA (K11R) displays fewer SIFs. **(a)** Images from HeLa cells infected with strains pSifA^WT^ and pSifA^SMUT^ compared for their ability to form SCVs (stained by anti-*Salmonella* LPS antibody) 4hours post infection. The integrity of SCVs were assessed by detecting the presence of the marker galectin-8 (scale 8 micron). **(b)** Co-localisation analysis for SCVs and Gal-8 in depicted conditions to assess the integrity of SCVs. **(c)** Image fields with cells infected with indicated strains stained by SIFs marker LAMP1 probed after 10 hours post infection (scale 50 micron). **(d)** Field with single cells displaying SIFs (LAMP1 staining) infected with indicated strains (sacle 10 micron). **(e)** Quantitative representation of percent of cells containing SIFs from samples infected by varied strains of *Salmonella*.

### SifA SUMOylation is necessary *Salmonella* replication and virulence

Virulence of bacterial pathogens is mainly dependent on their ability to replicate and disseminate inside their host. In preparation for our assays, the expression levels of both wild type and the mutant SifA were analyzed growing cultures of ΔSifA complemented with wild type and mutant SifA (SifA-K11R) respectively (fig. EV1a). Next, a basic growth curve conducted to assess the multiplication rate for all four strains SL1344, ΔSifA, pSifA^WT^ or pSifA^SMUT^ revealed these strains followed similar growth kinetics (fig EVI1b). Gentamycin protection assay (GPA) was performed in cultured cells infected with above mentioned four strains of *S.* Tm to investigate intracellular bacterial load in infected cells. Post infection, HCT-8 cells were monitored at 2, 7 and 16 hours revealed no change in colony forming units (CFU) at early time periods (2hpi) (fig 5a). However, at 7hpi, CFU from pSifA^SMUT^ infected cells was considerably lesser than other strains (fig 5a). The replication of ΔSifA at this point was very high as expected due to hyper replication of this strain owing to loss of vacuole (Knodler, 2015). After 16hpi, the difference in intracellular bacterial load from cells infected by both ΔSifA and pSifA^SMUT^ was significantly lower in comparison to their respective wild type controls (fig 5a). Aside from epithelial cells, we also carried out GPA in isolated primary bone marrow derived macrophages (BMDMs) from mice. *Ex vivo* infection of BMDMs by the above mentioned strains for a duration of 16hrs, yielded similar pattern where strain pSifA^SMUT^ replicated at a slower rate in comparison to pSifA^WT^ (fig 5b). Finally, the virulence potential of pSifA^WT^ and pSifA^SMUT^ were further tested in other cell lines HeLa (fig EV1c), CaCo2 (fig EV1d) and RAW 264.7 (fig EV1e) post 16 hours of infection. The pattern observed was analogous to previous findings obtained from HCT-8 and macrophages. Thus, it was evident that SifA^SMUT^ expressing strain exhibited a severely compromised replication *in vitro* in a range of different cell types.

**Fig. 5:**
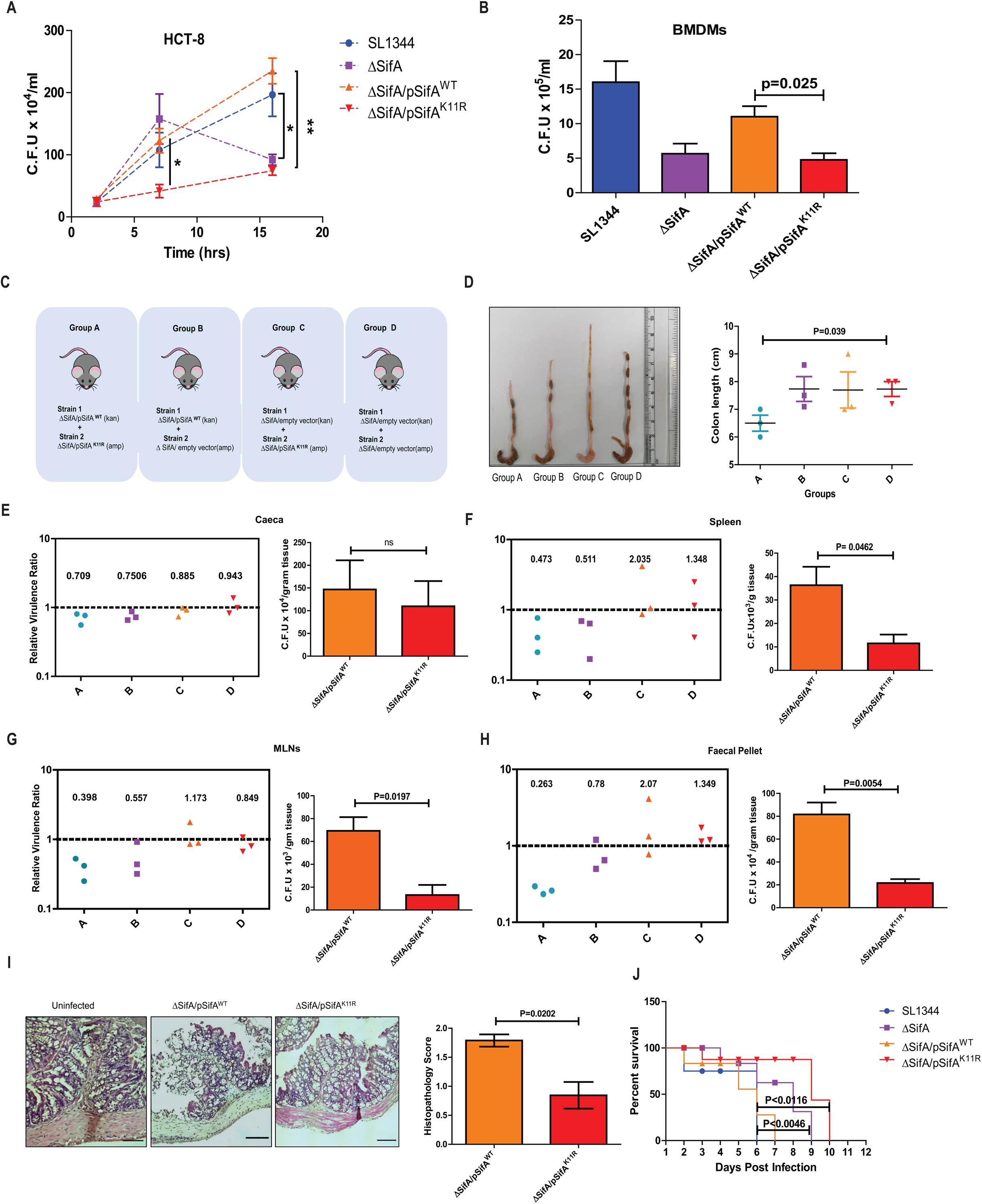
SifA SUMOylation is necessary for implementing virulence. **(a)** The schematic describing the mice groups infected with the indicated strains. CI assay involves infecting the animal with mixed inoculum consisting strain 1(WT) and strain 2 (SMUT) in equal ratio. **(b)** The colon morphology comparison from infected animals across the groups. **(c)** Graphical representation of colon length from shown groups with significant variation in group A (pSifA^WT^ + pSifA^SMUT^) and D (Vector control). **(d-g)** Relative virulence ratio calculated by the equation described in the text. CI with mean index is displayed for bacterial counts obtained from caeca, spleen, MLNs, and fecal pellet. The bar graph at the right compares the bacterial load plotted from mice individually infected by the indicated strains. **(h)** H and E studies showing increased intensity of inflammation in pSifA^WT^ infected mice proximal colon sections. The histopathology of the colon was quantified by considering parameters like epithelial damage, goblet cell loss and immune cell infiltration which were scored blind by a trained pathologist. **(i)** The survival curve representing no. of days lived by every infected mouse from all groups. Each drop signifies death of an animal in the group. Mice infected by ΔSifA and pSifA^SMUT^ strains showed prolonged survival owing to attenuated virulence.

**Fig.6: Schematic.**
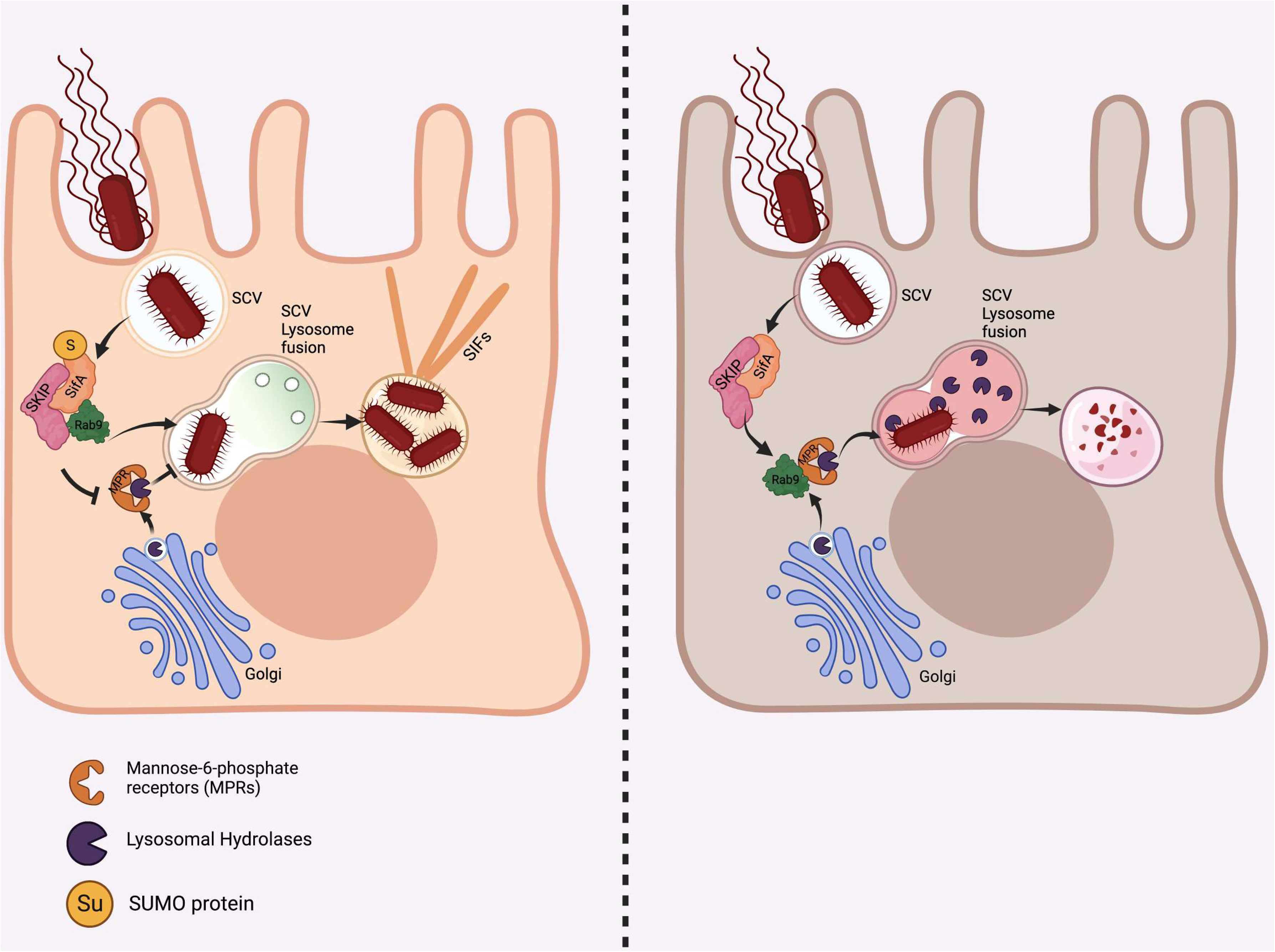
describing role of SifA SUMOylation in *Salmonella* pathogenesis. SifA SUMOylation is essential for Rab9 sequestration by PLEKHM2 which subverts CI-M6PR routing and thus protects SCV from lysosomal hydrolases. SifA^SMUT^ cannot alter this trafficking route which leads to SCV degradation. *This figure has been created using biorender software* (https://biorender.com/)

The attenuated replication of *S.* Tm bearing SifA^SMUT^ detected *in vitro prompted* us to examine the behavior of this strain in mice models of *Salmonella* infection. Streptomycin pre-treated C57BL/6 female mice model was adapted to carry out all the infections. A Competitive index (CI) assay was conducted to assess the fitness of strain carrying SifA^SMUT^ in comparison to those bearing SifA^WT^ (Macho *et al*., 2007). To actualize this assay, ΔSifA complemented with SifA^WT^ cloned in pET28a vector (vector 1, kanamycin selection) was mixed with SifA^K11R^ complemented in ΔSifA via pET21a vector (vector 2, ampicilin selection) in equal ratio to achieve a MOI of 5×10^7^ CFU/ml. The input inoculum contained equal volume of both cultures since they multiplied at same rate in LB media. Streptomycin treated mice post 24 hrs, were fed with the above stated mixed cultures by oral gavage, for a total duration of 48hrs after which animals were euthanized and tissue lysates plated for CFU enumeration. The panel (fig 5c) indicates the categorized mice groups (N=3) fed with the corresponding strains as shown. Colon harvested from these mice groups were assessed for their morphology, which exhibited maximum inflammatory features in group A (pSifA^WT^ + pSifA^SMUT^) whereas these features were minimally seen in group D (Vector1+Vector2) (fig 5d). Furthermore, the tissue homogenate from caeca, spleen, mesentric lymph nodes (MLNs), and fecal pellets were used to calculate CFUs on McConkey agar with appropriate selection markers (kan for WT and amp for SMUT). The following formula was used to calculate the Competitive Index:

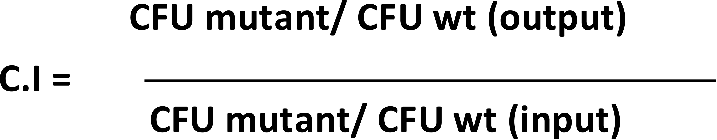

The CI signifying relative virulence ration calculated as mentioned above from organs are shown in fig 5 d-g. An index value close to 1 suggests both the strains replicate at equal rate, while a value farther from 1 propose one strain outcompetes the other. The mean index of CI values from three mice is also shown in the graph. As evident from these graphs, both pSifA^WT^ and pSifA^K11R^ strains were able to colonize and multiply similarly in mice gut like other groups. However, the index values for group A varied from group D significantly in other organs checked. Furthermore, the index values were comparable between groups A and B on the other hand groups C and D showed similar values meaning wild type strain outcompeted SMUT which replicated and disseminated identical to ΔSifA strain. To support the above data, CFU involving single infections (5×10^7^ CFU of individual strain per mice) performed in same organs under similar conditions, were also added beside their corresponding CI plots (fig 5e-h). Differences in bacterial load were largely observed in MLNs and faecal pellet. Histopathological analysis involving Hematoxylin and Eosin (H & E) studies done in proximal colon tissue sections from mice infected by indicated strains showed severity of inflammation varied between wildtype and SMUT (fig 5i). The histopathology score tested was higher for pSifA^WT^ compared to pSifA^K11R^. Finally, a survival assay was administered comprising SL1344 and ΔSifA in addition to complemented strains for evaluating their virulence in mice. Mice groups orally infected by depicted four strains (N=4) separately at the same time were left monitored to decide the progression of infection leading to bacteremia and death in these animals. The curve in fig 5j indicated the mice infected with complemented strain pSifA^K11R^ survived for more days (till day 10), like ΔSifA strain. However, mice groups infected by the wild type strains succumbed to infection much earlier. These data sets identified that SMUT SifA bearing strain is comparably attenuated in its virulence potential to spread Salmonellosis in animals.

Hereby, we concluded that SUMOylation of SifA during infection is essential for *Salmonella* to thrive in the host, overcoming lysosomal attack and its further dissemination.

## Discussion

The present study emphasizes a novel regulatory mechanism, where the effector SifA is seen to undergo modified by SUMOylation, a host PTM mechanism, for the progression of Salmonellosis. SPI-2 effector SifA is required for dissemination of *Salmonella* and establishing systemic infection in mice (Brumell, Rosenberger, *et al*., 2001; Fierer *et al*., 2012). At the cellular level, SifA is responsible for SCV maintenance and SIF generation, two phenotypic hallmarks associated with the virulence of *Salmonella* (Beuzón *et al*., 2000; Brumell, Goosney and Finlay, 2002). Mechanistically, the presence of SifA subverts trafficking of Rab9-M6PR mediated lysosomal hydrolases targeting SCVs, thereby protecting it from lysosomal degradation (McGourty *et al*., 2012). To perform these numerous functions, SifA interacts directly or indirectly with multiple host adaptors including SKIP, PLEKHM1, Rab7, Rab9, LAMP1 simultaneously associating with few other SPI-2 effectors (Ohlson *et al*., 2008; Diacovich *et al*., 2009b; McEwan *et al*., 2015; Sindhwani *et al*., 2017). In this given scenario, the imperative question that needs to be addressed is the factors regulating such strategic placements of these proteins within the complex network. PTM mechanisms are tools regulating almost entire cellular events, which has been shown to be targeted by pathogenic effectors to steer control of host cellular physiology. As reported earlier, SifA undergoes PTMs prenylation and S-acylation for membrane anchorage and achieve destined localization in infected cells (Reinicke *et al*., 2005). Our group reported PTM by SUMOylation to be drastically altered upon *Salmonella* infection (Verma *et al*., 2015). Besides, it was demonstrated that the intracellular fate of *Salmonella* is significantly impacted by the SUMOylation of GTPase Rab7 (Mohapatra *et al*., 2019). In this study we showed that Salmonella controls the SCV stability by not allowing Rab7 to undergo SUMOylation. However, it was not clear if any of the other protein which are in the vicinity of Ran7, such as SifA, SKIP and others undergo SUMOylation. Other components of vesicular transport pathway have been shown to undergo SUMOylation, like Rab17 undergoes SUMOylation which directs its interaction with syntaxin 2 (Striz and Tuma, 2016). Several components of autophagy have been shown to to be regulated by SUMOylation. This includes Vps34 SUMOylation required for autophagosome formation <u>(Yang *et al*., 2013)</u>. Lysosome mediate clearance of Huntington’s disease protein Huntingtin relies of SUMOylation (Thompson *et al*., 2009). Similarly, trnaslocation of RhoB, a Rho GTP binding family protein, to lysosomes depends on its SUMOylation (Liu *et al*., 2018).

Further, other relating studies have shown effectors of various pathogens e.g., AmpA from *Anaplasma* (Beyer *et al*., 2015) and TRP120 from *E. chaffeensis* (Dunphy, Luo and McBride, 2014) getting SUMOylated during infection for aiding the pathogen survival inside hosts. Hence, the strong background from all these studies and a need for convincing hypothesis to provide insights regarding functional regulation of SifA paved the foundation of this study.

The current study identified SifA undergoing SUMOylation at its 11^th^ lysine, and further proved how *Salmonella* exploited this modification for its benefit.

It is well known that through its interaction with SKIP, SifA contribute to the regulation kinesin-1 mediated anterograde movement of late endosomal compartments (Dumont *et al*., 2010). Therefore, it can be concluded that cytoskeletal modulations brought up by bacterial effectors are the fundamental physical processes that work up for creating a niche for pathogen. The functions of SUMOylation are well explored in processes like transcriptional regulation, DNA repair, sub cellular localization, substrate folding etc., (Sarangi and Zhao, 2015; Abe *et al*., 2017; Rosonina *et al*., 2017; Yang *et al*., 2017) nonetheless their role in regulation of cytoskeletal elements is currently surfacing. Recent studies have revealed that actin gets SUMOylated for its nuclear transport (Hofmann *et al*., 2009), also alpha tubulin undergoes SUMO modification to exert control over microtubule assembly (Feng *et al*., 2021). One important aspect associated with control of cytoskeletal proteins is cellular trafficking. SUMOylation enable their substrate cargoes to be transported in both anterograde and retrograde directions reversibly. A study from literature concerning RNA binding La chaperone protein demonstrated that directionality of its transport is dependent on its SUMOylation at lysine 41 in neuronal axons (van Niekerk *et al*., 2007).

Hence, we presume that SifA SUMOylation provides a scaffold to carry out various cellular trafficking that decide the fate of bacterial existence in host cells. Misrouting Rab9 conjugated M6PR vesicles preventing SCV acidification, (McGourty *et al*., 2012), utilizing PLEKHM1 Rab7 complex for mobilizing phagolysosomal membranes for SCV generation, (McEwan *et al*., 2015; Sindhwani *et al*., 2017) binding to SKIP Kinesin-1 complex for SIF extension (Ohlson *et al*., 2008) etc are few of the cellular transport orchestrated by SifA. SifA SUMOylation supposedly impart control to these complex trafficking events during pathogenesis. The host SUMO ligases and deSUMOylases involved in maintenance of SifA SUMOylation and the spatiotemporal regulation of above stated events directed by SUMO are beyond the scope of this work. Nevertheless, our work indicates SUMO is integral component of functional SifA and hampering it collapses the prominent phagolysosomal alterations brought by SifA during pathogenesis. Moreover, the present work cites the first evidence of a *Salmonella* effector undergoing SUMOylation for its sustenance inside hosts and disclose additional layers of ramifications taking part in host pathogen cross-talk.

## Materials and Methods

### Cell culture and treatments

Human adenocarcinoma Cell lines HCT-8, were cultured in Rosewell Park Memorial Institute (RPMI) media supplemented with 14 mM NaHCO3 (Sigma), 15 mM HEPES buffer (pH 7.4),1 mM sodium pyruvate (GIBCO), 40 mg/L penicillin (GIBCO), 90 mg/L streptomycin (GIBCO), and 10% foetal bovine serum (FBS) (GIBCO). Human colorectal adenocarcinoma cell line CaCo2 and human cervical carcinoma cell line HeLa, were grown in Dulbecco’s modified Eagle’s medium (DMEM) comprising14 mM NaHCO3, 15 mM HEPES buffer (pH 7.5), 40 mg/l penicillin, 90 mg/l streptomycin, and 10% FBS. Bone Marrow Derived Macrophages (BMDMs) were prepared from the femur of C57BL/6 mice by flushing them aseptically in DMEM. Cells thus obtained were first treated with RBC lysis buffer (Sigma) and then cultured and differentiated in DMEM supplemented with 10 mM HEPES, non-essential amino acids (GIBCO), 10% FBS and 20% L929 conditioned media. The pharmacological inhibitors included in this study includes cycloheximide used at a concentration of till the indicated time, proteasomal inhibitor MG132 at 20µM working concentration for a total duration of 4hours. Vacuolar ATPase inhibitor bafilomycin A1 treatment was given at a concentration of 100nM for total duration of 4 hours.

### Bacterial Strains, Plasmids, and *In vitro* infection

*Salmonella* Typhimurium strain SL1344 was used throughout the studies. The SifA deletion knockout strain (ΔSifA) in SL1344 background together with the parent strain were kindly gifted by Beth McCormick, University of Massachusetts Medical School, MA. The coding sequence of SifA amplified from SL1344 genome were cloned in a series of prokaryotic expression vectors including pBH-HA from Roche, pET28a, pET21a and pGEX-6p1 vectors for various experiments. SUMO intolerant SifA (SifA SMUT) was generated via site directed mutagenesis (SDM) in all these above vectors. Plasmids bearing wild type and mutant clones were transformed into the ΔSifA strain via electroporation for the generation of complemented strains ΔSifA/pSifA^WT^ and ΔSifA/pSifA^K11R^ respectively. All the mentioned bacterial strains were cultured in Luria Bertani (LB) broth at 37°C aerobically for 8 h serving as the primary culture. A secondary culture grown by inoculating the primary culture in 12 ml LB broth in standing white cap tubes under stationary hypoxic conditions overnight at 37°C were used to infect epithelial cells HCT-8, HeLa and Caco2 at a multiplicity of infection (MOI) of 1:40. BMDMs were infected similarly at a MOI of 1:10. Mammalian vector bearing SifA cloned in peGFP-C1 vector kindly gifted by Dr. Amit Tuli, IMTECH, Chandigarh, India was used for overexpression studies.

### Cell Transfection

Transfection studies were conducted in HCT-8 and HeLa cells by the standard transfection protocol employing reagents lipofectamine 2000 and lipofectamine LTX (Invitrogen, USA) respectively. Day prior to transfection, 2.5×105 cells were plated in 24-well plates to obtain 80% confluency. Briefly, 1µg of required plasmid and the transfectant reagent taken at a ratio of 1µgplasmid: 2µl of reagent were separately diluted in minimum serum media Opti-MEM (Invitrogen, USA) and incubated for 5 min. The two mixtures were then mixed and incubated for another 15 min at room temperature and subsequently supplied to cells growing in Opti-MEM followed by an incubation of 24 hours, after which cells were replenished with complete media.

### Yeast Two Hybrid Assay

The bait fusion construct in vector pGBKT7 expressing GAL4 DNA binding domain (BD) and prey fusion construct in vector pGADT7 expressing GAL4 activation domain (AD) were co-transformed in yeast strain (Y187, Clonetech’s matchmaker Gold Y2H system). Transformants grown on SD media plates devoid of auxotrophic markers leucine and tryptophan at 30°C were replated on selection plate further lacking histidine and adenine to monitor bait prey interaction.

### Protein Purification and *In vitro* SUMOylation Assay

*E. coli* expression vector pGEX6P1 encoding SifA^WT^ or SifA^K11R^ were transformed in strain BL21 followed by IPTG induction at a concentration 0.05mM at 37◦C. The fusion construct GST SifA^WT^ or GST SifA^K11R^ were purified by using glutathione sepharose beads (GE Healthcare) and further cleaved with thrombin for tag separation. Proteins further purified by passing in a superdox 200 column were subjected to SUMO conjugation reactions with the help of commercially available *in vitro* SUMOylation assay kit (Enzo Life Sciences). Kit components comprising SUMO machinery enzymes and ATP were added to 500 nM of wild type SifA and SMUT SifA proteins separately in a 20 µl reaction mix along with provided controls at 37°C for 1 hour as per manufacturer’s instructions. The reactions were stopped and processed by adding Laemmle buffer after which it was loaded onto SDS-PAGE and immunoblotted to be probed by SUMO-1 and SUMO2 antibodies.

### In-bacto SUMO Conjugation Assay

This assay again included co-expressing SifA^WT^ or SifA ^SMUT^ from pGEX6P1 vector in *E. coli* BL21 along with plasmids encoding SUMO machinery enzymes pE1E2S1 (SUMO-1) or pE1E2S2 (SUMO-2) for the overproduction of SUMO conjugated substrates in bacteria (Uchimura *et al*., 2004). Rab7 was chosen as positive control for this assay (Mohapatra *et al*., 2019). Double transformants growing on plates supplied with ampicillin and chloramphenicol were cultured in LB in presence of 0.05mM IPTG for induction at 37◦C for 3 hours which is followed by lysate preparation and analysis by western blot.

### Immunoblotting

Cells lysates were prepared in Radioimmunoprecipitation assay buffer (RIPA) (Sigma) supplemented with Protease Inhibitor Cocktail (G biosciences). These lysates were further denatured by boiling in 2x SDS Laemelli buffer (20 mM Tris-HCl pH 8.0, 150 mM KCl, 10% glycerol, 5 mM MgCl2 and 0.1% NP40) at 95◦C for 10 minutes. CBX™ protein assay kit (G-Bioscience, USA) was utilized for protein quantification. Lysates were then separated by routine sodium dodecyl sulfate–polyacrylamide gel electrophoresis (SDS-PAGE) in tris glycine running buffer. Resolved gels were transferred on to nitrocellulose membrane (BioRad) after which the blots were probed with antibodies against SUMO-1 (CST), SUMO-2 (CST), GST (Sigma), eGFP (Abcam), PLEKHM2 (Abcam), CI-M6PR (Abcam) Actin (Thermo Scientific), and GAPDH (Invitrogen). The instrument Image Quant LAS4000 from GE was used for blot developing and processing and densitometry analysis was done using the Image J software.

### Immunoprecipitation

For immunoprecipitation HCT-8 cells were lysed in non-denaturation immunoprecipitation lysis buffer (Pierce, Thermo Scientific) supplemented with 20 mM N-Ethyl Maleimide (NEM), (Sigma) and 2x protease inhibitor cocktail (G biosciences) for 20minutes on incubation on ice. Lysates were prepared by centrifugation at 15700 g for 10 min at 4°C and collecting the supernatant. The lysates obtained were subjected to preclearing by incubation with protein G sepharose beads (GE) for 30 minutes at 4°C on an end-to-end rotor. Precleared lysates after centrifugation were then incubated with anti-SUMO1 antibody at 4°C on an end-to-end rotor for overnight. Next day, the incubated lysates were precipitated by addition of protein G sepharose beads for 3hours at 4^◦^C. Precipitations were also carried by IgG raised in same host (Merck Millipore) as the isotype control. The antigen-antibody complex obtained after washing with lysis buffer were processed in Laemmle buffer for immunoblotting.

### Immunofluorescence

HeLa and HCT-8 cells were grown on cover slips embedded in 24-well plates. In assays monitoring lysosomal activity ratiometric probes like DQ-BSA, lysotracker and lysosensor were incorporated in cells prior to fixation as per manufacturer’s instructions. Post infection, cells were washed thrice in 1× phosphate buffered saline (PBS) and fixed in 4 % methanol free paraformaldehyde (CST) for 15 min at room temperature (RT). Following washing, cells were permeabilized in 0.5% saponin in PBS for 10 minutes at RT after which they were blocked in blocking solution (1%BSA in PBS containing 0.1% Tween 20) for 1 hour at RT. Cells were incubated in primary antibodies against LAMP1 (Sigma), Rab9 (Invitrogen), GFP (1:200), Galectin-8 (Abcam), CI-M6PR (Abcam), SUMO-1(Invitrogen), *Salmonella* LPS (Abcam) prepared in blocking solution for overnight at 4^◦^C. Next days, cells were washed four times in PBS and proceeded for incubation in fluorophore conjugated secondary antibodies for 2hrs at RT in PBS containing 0.5% BSA and 0.1% Tween-20. Cells were finally washed three times and incubated in 4,6-diamidino-2-phenylindole (DAPI) (1 µg/ml; Sigma-Aldrich) for 5 min in dark. Mounting was done using prolong diamond antifade (Invitrogen). Images were acquired in Leica SP8 confocal microscopy under 63× oil immersion objective. Colocalization analysis and intensity calculations were obtained from acquired images using Image J software.

### Proximity Ligation Assay

Proximity Ligation Assay (PLA) was conducted using Duo-link PLA kit (Sigma) as per manufacturer’s protocol. Cells after permeabilization as described before were blocked in PLA blocking buffer followed by incubation in primary antibody against GFP (rabbit) and SUMO-1(mouse) overnight. Next day, the coverslips were washed in buffer A provided in the kit. Further, the samples were incubated with plus and minus end PLA probes supplied by the manufacturer. Hereafter the attached probes were ligated and subsequently PCR amplified for 100 minutes using the provided reagents. All the reactions were performed at 37◦C in a humidity chamber. Finally, the samples were washed in buffer B and mounted onto a slide with the supplied mountant containing DAPI for visualization. The PLA signal punctae were quantified from indicated no. of cells across each sample and plotted using the graph pad software.

### Gentamycin Protection Assay

Infection in cells by various strains at an MOI of 1:40 in medium lacking pen strep were allowed to proceed for 1 hr at 37°C in the incubator, followed by treatment with antibiotic gentamicin (Gibco) at a higher concentration of 100ug/ml in media for 1 hr to eradicate extracellular bacteria. After this step, cells were washed in sterile PBS followed by further incubated in the media containing gentamicin at a concentration of 20 µg/ml for the rest of the infection. At the end, infected cells were lysed using PBS containing 0.1% triton X-100 detergent followed by dilution plating on LB plates added with streptomycin (50 µg/mL). The plates were incubated in a 37^◦^C incubator and individual CFU were counted and graphically represented.

### Mice Infections

All in vivo studies were conducted in C57BL/6 female mice (6–8 weeks). All the animal experiments were carried out in the Small Animal Facility of Regional Centre for Biotechnology (RCB). The experiments were performed after approval by the RCB Institutional Animal Ethics Committee (approval no. RCB/IAEC/2017/019 & RCB/IAEC/2021/087). For the induction of colitis streptomycin pre-treated colitis model was employed (Barthel *et al*., 2003). Food and water were removed 4 hrs prior to infection followed by a treatment with 20 mg/Kg of streptomycin by oral gavage for the clearance of microbiota. After 24 hours, *Salmonella* SL1344 strain at a MOI of 5×10^7^ CFU per mice was fed using oral gavage after 4hrs of withdrawal of food and water. Post 48 hours of infection, mice were euthanized and various organs including colon, spleen, MLNs, and faecal pellets were harvested. The tissues from organs were homogenized in PBS+ 0.1% Triton-X solution in Precelly’s bead beater followed by dilution plating on Mc Conkey agar plates containing streptomycin (100µg/ml) for bacterial enumeration. For the survival assay, mice infected by the mentioned strains as described above were left monitored for death succumbing to progression of infection. A survival curve indicating the no. of days each mouse survived across all groups were plotted using graph pad prism software.

### Competitive Index Assay

This assay involves infecting the same mice with mixed inocula comprising both wild type and mutant protein expressing strains in equal ratio (Auerbuch, Lenz and Portnoy, 2001; Macho *et al*., 2007). The strains ΔSifA/pSifA^WT^ (strain1) ΔSifA/pSifA^K11R^ (strain 2) were cloned in pET28a vector (Kan resistance) and pET21a vector (Amp resistance) respectively, thus enabling a plasmid-based antibiotic marker differentiation. The input bacterial inoculum containing equal concentrations of both strain 1 and strain 2 totaling to a MOI of 5×10^7^ CFU standardized by absorbance based CFU determination were fed to the C57BL/6 mice as described earlier. Homogenized tissue lysates from organs collected post 48hrs of infection were simultaneously plated on separate McConkey agar strep plates supplied with kanamycin (50µg/ml) or ampicillin (100µg/ml). CI was calculated as the CFU of mutant-to-wild-type ratio obtained from target sample, divided by the corresponding CFU ratio in the inoculum (input sample).

### Haematoxylin and Eosin Staining

Proximal colon sections were fixed in 10% formalin buffer for two days at room temperature and frozen in holders with a thick layer of cryomatrix (Thermo Scientific) and stored at −80◦C till use. Five micrometre thick sections were cut onto glass slides in a microtome and processed for haematoxylin (Sigma) and eosin (Sigma) staining. The slides and mounted using DPX mountant (Sigma) and bright field images were taken using a Nikon (NY, USA) inverted fluorescence microscope.

### Statistics

The results are expressed as the mean standard error from an individual experiment done in triplicate. Data were analysed either with one way ANOVA followed by standard unpaired two-tailed Student’s t test, with *p* values of .05–.001 considered statistically significant. All graphs and stats were plotted using the graph pad prism software.

## Acknowledgements

We thank the RCB Central Instrumentation Facility (CIF) and NCR Cluster Small Animal Facility. We are thankful to Prof. Sudhsnshu Vrati for supplying reagents for Proximal Ligation Assay. This work was supported by STARS grant (STARS/APR2019/BS/730/FS) and to C.V.S. and core funding from Regional Centre for Biotechnology, Faridabad, India.

## Conflict of Interests

WE DECLARE THAT THERE IS NO CONFLICT OF INTEREST AMONG ANY OF THE AUTHORS OF THIS WORK.

| Sl. No. | Author Name | Contributions |
| --- | --- | --- |
| 1. | Ms. Hridya Chandrasekhar | Fig 1f, 1h, 1i<br>Fig 2 (a-j)<br>Fig 3 (a-h)<br>Fig 4a, 4b |
|  |  | Fig Ev1a, Ev1b, Ev1d<br>Fig 5c to 5i |
| 2. | Dr. Gayatree Mohapatra | Fig. 1c, 1d, 1e, 1g<br>Fig. 4c, 4d, 4e<br>Fig. Ev1c, Ev1e<br>Fig. 5a, 5b |
| 3. | Dr. Mukesh Singh | Fig. 5j, Survival Curve |
| 4. | Dr. Sarika Rana | Prepared all clones and mutants via SDM |
| 5. | Dr. Navneet Kaur | Performed protein purification for figures<br>1c, 1d and 1e |
| 6. | Ms. Sheetal Sharma | Fig 1a, Y2H assay |
| 7. | Dr. Amit Tuli | Fig 1a, Y2H assay |
| 8. | Dr. Prasenjit Das | Pathologist, performed the histopath scoring<br>for proximal colon sections in fig 5h |
| 9. | Dr. C.V Srikanth | Corresponding Author: Idea &<br>conceptualization, analysis and<br>interpretation |

---

**Expanded View Figure 1.**
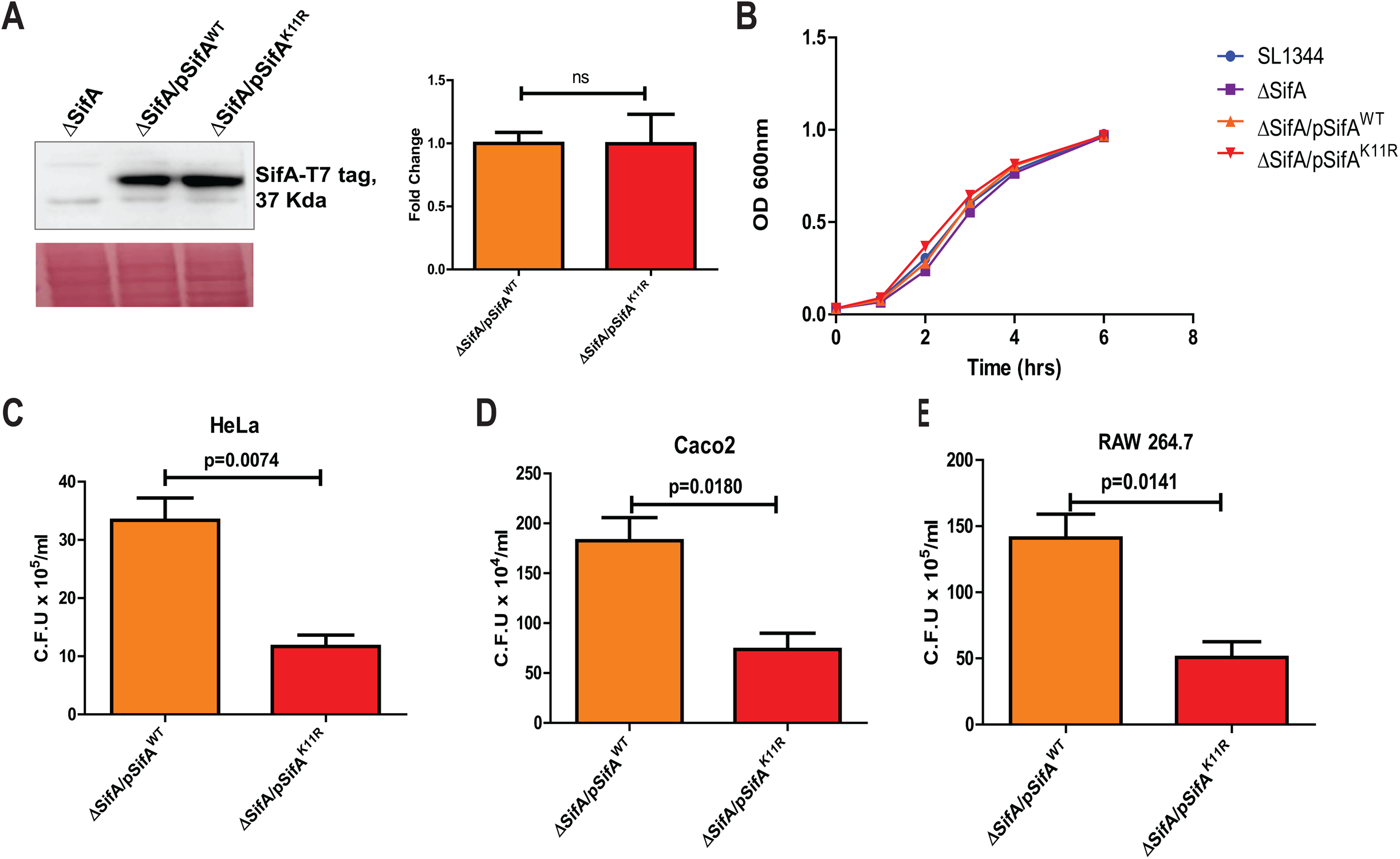
**(a)** The expression levels of SifA^WT^ and SifA ^SMUT^ proteins from ΔSifA complemented with SifA^WT^ (pSifA^WT^) and SifA ^SMUT^ (p SifA ^SMUT^) respectively. (**b**)The growth curve indicating division rate of strains SL1344, ΔSifA, pSifA^WT^ and pSifA ^K11R^. (**c**) Gentamycin protection assay indicating bacterial burden in HCT-8 cells infected by the shown strains at 2, 7 and 16 hours post infection (* denotes p value of 0.0215 and ** equals p= 0.0018). **(d)** Intracellular bacterial load from isolated primary BMDMs infected by shown strains 16hpi. Comparison of CFU results obtained from pSifA^WT^ and pSifA ^KIIR^ infected **(e)** Hela (f) Caco2 cells and (g) RAW 264.7 cells post 16 hours.

